# Career Self-Efficacy Disparities in Underrepresented Biomedical Scientist Trainees

**DOI:** 10.1101/2022.10.21.512368

**Authors:** Deepshikha Chatterjee, Gabrielle A. Jacob, Susi Sturzenegger Varvayanis, Inge Wefes, Roger Chalkley, Ana T. Nogueira, Cynthia N. Fuhrmann, Janani Varadarajan, Nisan M. Hubbard, Christiann H. Gaines, Rebekah L. Layton, Sunita Chaudhary

## Abstract

The present study examines racial, ethnic, and gender disparities in career self-efficacy amongst 6077 US citizens and US naturalized graduate and postdoctoral trainees. Respondents from biomedical fields completed surveys administered by the National Institutes of Health Broadening Experiences in Scientific Training (NIH BEST) programs across 17 US institutional sites. Graduate and postdoctoral demographic and survey response data were examined to evaluate the impact of intersectional identities on trainee career self-efficacy. The study hypothesized that race, ethnicity and gender, and the relations between these identities, would impact trainee career self-efficacy. The analysis demonstrated that racial and ethnic group, gender, specific career interests (academic principal investigator vs. other careers), and seniority (junior vs. senior trainee level) were, to various degrees, all associated with trainee career self-efficacy and the effects were consistent across graduate and postdoctoral respondents. Implications for differing levels of self-efficacy are discussed, including factors and events during training that may contribute to (or undermine) career self-efficacy. The importance of mentorship for building research and career self-efficacy of trainees is discussed, especially with respect to those identifying as women and belonging to racial/ethnic populations underrepresented in biomedical sciences. The results underscore the need for change in the biomedical academic research community in order to retain a diverse biomedical workforce.

## Introduction

Systemic barriers preventing racially and ethnically diverse populations from inclusion in biomedical research have been a long-standing issue (1–4). However, the existence of “systematic and intentional” exclusion of people based on their ethnicity or race (5) has been acknowledged explicitly only in recent years. Despite some progress, gender, race, and ethnicity continue to pose barriers to participation in biomedical higher education and to progression to academic leadership in the US (2, 6). While increased attention has been paid to addressing gender disparities in the pre-and postdoctoral arena, particularly in the biomedical sciences, race, and specifically the intersection of gender and race, continue to hinder the advancement of such trainees, particularly at the higher academic ranks (7,8). Studies of intersectional identities that focus on the cumulative effect of belonging to multiple social groups that are underrepresented (e.g., being Black *and* female) have shown that for trainees with intersectional identities these disparities may be compounded, and therefore further efforts are needed to remove these barriers. (9). In line with the literature on intersectionality, the current study defines intersectional identity as belonging to multiple social identity groups simultaneously, such as gender and race (10, 11). Hereafter, we refer to individuals from racial/ethnic and gender minorities as underrepresented (UR) trainees. To be clear, systemic barriers are at the root of the differences we hypothesize exist, and the various impacts measured across any particular group or combination of identities represent the cumulative (or interactive) effects of the lived experiences of trainees with particular social identities (12).

Scholars contend that interest in science, technology, engineering, mathematics (STEM; see Supplemental Information S1 for list of acronyms) is developed over the years through socialization and educational experiences (e.g., starting from K-12 experiences, through undergraduate, and graduate education), and that a lack of participation of racial/ethnic and gender minorities in STEM careers starts early (13–15). For example, in a longitudinal study that tracked students’ transition from high school to postsecondary careers, higher math self-efficacy during high school increased the likelihood of choosing STEM fields for White students but not for students from underrepresented groups (15) (note that White is capitalized throughout to indicate that it is a racial category as well, and to decenter Whiteness; (12)). Such sorting into different professions exemplifies the early loss of students from underrepresented backgrounds at various points along their education and career advancement, resulting in relatively few URs holding senior-level positions in STEM. Relatedly, individuals identifying with racially underrepresented groups are less likely to choose STEM fields of study, further compounding the problem of reduced representation and retention, once they embark on an academic research path.

Career choices can be studied through the lens of social cognitive career theory (SCCT; (16), which describes how individuals gain interest in specific careers, choose from the array of career options available to them, and engage in career-relevant activities to ensure career success. As described in SCCT, career self-efficacy (CSE) is the confidence to take ownership of one’s career plans and outcome expectations. Furthermore, according to SCCT, personal variables such as gender and race (among others) can impact the selection of, and access to the types of training opportunities that might be available. Restricted opportunities and unwelcoming environments in STEM can impact an individual’s interest, career choices, self-efficacy, and expectations regarding possible STEM career outcomes (16–18).

Gender and racial disparities have been noted on all academic levels (19). Once individuals start their careers as tenure-track faculty, issues of gender and racial biases (conscious and unconscious) continue to impact their progress (9, 20, 21). Self-efficacy is one essential factor affecting persistence in academia (22). Hence, it is important to study whether the CSE of trainees varies as a function of their gender and race. To the authors’ knowledge, studies on the role of intersectional identities, i.e., the multiplicative role of race and gender related to CSE in the biomedical sciences, are still uncommon (exceptions include (18, 23, 24)). Also noteworthy are recent calls to study how those who belong to two or more underrepresented social groups (for example, being Black *and* female) fare in the biomedical fields (9,21).

The present study, which evolved as a collaborative effort among institutions funded by the NIH BEST program (25), makes two significant practical and theoretical contributions. First, it addresses effects of race and gender on CSE of graduate students and postdoctoral fellows from racial and ethnic groups that are underrepresented (UR) versus well-represented (WR) in the biomedical sciences. Given the need to harness the STEM workforce for the US to maintain an internationally competitive position, it is critical that individuals who identify with UR racial and gender groups have a fair and equitable chance to participate. Several studies have shown that equity is missing in many biomedical fields (2, 4, 23), but such studies often are often limited to a small sample size. While these studies provide valuable insights, they may be missing effects that would be detectable with a larger sample size, thereby preventing meaningful policy inferences and actions. For example, Gibbs and colleagues’ (2) analysis of career interest patterns in biomedical science doctoral students included a total of 1500 respondents. Although this number seems a sizeable sample, of the 1500 respondents, only 5.8% men (n=87) and 12.6 % women (n=189), or 276 respondents total, self-identified as members of underrepresented racial/ethnic groups. Similarly, Layton and colleagues (4) had also recruited a large national sample of 8099 respondents (3669 usable data), but again, only 7% of those self-identified as members of from underrepresented groups (n=225; 81 men, 144 women). Low representation has led to various studies reporting conflicting results regarding the impact of gender and race on CSE, including how CSE affects career choice and the job search process. For example, while some analyses found that gender and race impact outcomes negatively, some found positive impacts, and others found no effect (2, 18, 23, 26, 27). Trainees who completed a common “entrance survey” at NIH BEST-funded institutional sites were analyzed; this more comprehensive survey data from historically UR gender and racial/ethnic groups yield important insights for practitioners, educators, policymakers, and sponsors.

While the bulk of the previous research on SCCT—particularly the work on CSE— assesses the impact of gender and race separately, the current study employs an intersectional perspective and based on prior research, introduces two main hypotheses. 1) The double jeopardy hypothesis (11) states that holding two minority identities can exacerbate negative outcomes; this hypothesis has gained some support in the literature (28–32). 2) A second competing hypothesis suggests a multiple identity advantage (10) whereby occupying double-minority identities can buffer an individual against negative outcomes related to a singular minority identity.

Understanding the role of intersectional identities is vital for improving support for underrepresented trainees and working to remove systemic barriers. The current work explores how individuals with intersectional identities and training in the biomedical sciences rate their CSE. If some individuals with intersecting, underrepresented racial, ethnic, and/or gender identities have to bear a multiple jeopardy effect, this result would be critical to keep in mind when addressing systemic barriers to mitigate negative outcomes. If, however, for some individuals, intersectionality provides a buffering effect, then Principal Investigators (PI) and policy makers could reframe the problem to tailor more nuanced solutions to meet the needs of specific populations. In either case, it will be essential to consider how issues are framed because research on achievement in STEM fields shows that solely addressing the impacts of exclusion of underrepresented individuals without acting on the systemic inequities can itself create and sustain negative outcomes (33–35).

In addition to systemic barriers for underrepresented groups, a lack of career preparation opportunities across graduate and postdoctoral training was identified in the Biomedical Workforce Working Group Report (36), which called for a substantial change in doctoral career development, prompting the National Institutes of Health to fund the Broadening Experiences in Scientific Training initiative (NIH BEST; (25)). The NIH BEST consortium was founded to provide better exposure of biomedical graduate and postdoctoral trainees (hereafter referred to as trainees) to the vast array of career fields in the biomedical sciences. This consortium of 17 US universities recognized that targeted career development training also had the potential to address inequities related to the exclusion of UR minority trainees and they collected standardized demographic and survey data across all participating institutions. The study presented here relied upon this robust national source (25). Analyses were confined to trainees who are US citizens or naturalized citizens. International trainees (defined as any classification of temporary visa holders or green card holders) were excluded from the present analysis as they may have different socio-cultural expectations and may encounter additional unique barriers to advancement in graduate and postgraduate training (37). This international trainee group deserves separate analysis regarding career decision-making and will be the subject of future work.

The current study aimed to document the self-reported CSE of trainees from underrepresented (UR) groups compared to BEST trainees from well-represented (WR) groups. In addition, to aid biomedical policymakers and academic administrators in their efforts to provide equitable opportunities and enhanced inclusiveness for all, the study analyzed the impact of the trainees’ career interest and seniority in training on self-efficacy. The NIH BEST program aimed to “seek out, identify and support bold and innovative approaches designed to broaden graduate and postdoctoral training” (38), included a strong interest to understand how individuals from underrepresented groups feel about pursuing careers in the biomedical field after completing their scientific training (39,40).

### Career self-efficacy: Role of Gender and Race

Lent and Hackett’s (41) SCCT extended Bandura’s (42) Social Cognitive Theory by positing that in career contexts, both CSE and career-related outcome expectations are vital components of success, and that both are influenced by demographics, such as gender and race as well as other personal factors like socioeconomic status and prior learning experiences (41,42). In SCCT, CSE is a construct that encompasses one’s belief in one’s own capacity to execute behaviors in pursuit of one’s career goals and is defined by Lent and Hackett (41) as “judgments of personal efficacy in relation to the wide range of behavior involved in career choice and adjustment” (p. 349) (41). CSE guides an individual’s choice of careers, how people engage with their chosen career tracks, and even impact adjustment processes they use in navigating career contexts. Career outcome expectations are beliefs that one’s successful engagement in certain career-related behaviors will yield desirable career outcomes. Individuals who are career self-efficacious (versus those who are not) will make positive (versus negative) cognitive appraisals of their future performance capabilities in a career domain (43). For example, career self-efficacious people will report feeling confident in pursuing their desired career paths and career goals. They will also report confidence in networking and seeking career advice from relevant stakeholders. In addition, they are likely to report confidence in figuring out how to best achieve their career goals (42-44).

The SCCT holds that personal factors like gender and race can become facilitators and/or impediments of success depending on an individual’s perception of compatibility with different academic/work environments. These personal factors engage with contextual factors of academic and/or work contexts (e.g., mentoring support, exposure to different role models, barriers such as financial support available, and family support), and together these forces create conditions that guide career interests, expectations, and career beliefs and behaviors (45). Gender and race can seem to restrict the type of career opportunities that trainees believe are accessible to them (46), and along with family and cultural expectations (47,48), they can disproportionately burden individuals from underrepresented groups with tokenism, stereotyping, and implicit bias in academic and professional domains.

From an early age, socialization differences yield differential patterns of career engagement for men versus women and individuals from well-represented versus underrepresented groups (49). Dewsbury and colleagues (50) studied how first-generation individuals from underrepresented groups in biological sciences found that in addition to engaging in an arbitrary career search process, racially underrepresented students (a) prioritize economic gains over intellectual fulfillment in a career, (b) seek to integrate the pressure of familial and cultural expectations to contribute to their family’s well-being versus devoting themselves primarily to a demanding career of their own interest, and (c) worry about being seen as token minorities in STEM careers (50). Gibbs and colleagues (2) found that underrepresented minorities of all genders were less likely to choose careers in research-intensive roles than well-represented men in biomedical sciences (2). African American and Asian doctoral biomedical trainees chose non-academic careers for financial rewards while White and Hispanic trainees chose non-academic careers to avoid the pressures of seeking research funding; furthermore, while women chose non-academic jobs to minimize pressures of seeking research funding, men chose non-academic jobs for higher financial rewards (51).

Beyond objective barriers such as financial rewards and research funding, concerns about numerical representation or “tokenism” create psychological threats and discomfort to minorities (52). In addition, such concerns can create and perpetuate structural barriers. For example, in academic and work contexts (referred to hereafter as performance contexts) that are male-dominated (e.g., engineering), White men benefit from the central and influential positionality in the organizational network of similar others, thereby disadvantaging women (53-56). Similarly, performance contexts can also be racialized as White-typical and penalize people from underrepresented groups (57,58); indeed, most organizations in the United States are White-majority (59). In fact, individuals from underrepresented groups report lower networking gains as compared to White trainees (24). Furthermore, underrepresented group members often conceive themselves as delegitimized as “diversity quota” or “affirmative action” hires (60). Indeed, underrepresented groups often face barriers and toxic climates that contribute to low advancement rates and/or even high attrition rates (21, 61). For example, Sheltzer and Smith (14) found that male faculty were less likely to hire female graduate students and female postdoctoral trainees, and this problem was compounded when “elite” male faculty made the hiring decisions (14).

The awareness that others may see them in stereotypical ways can be detrimental to confidence and self-esteem (35, 52,62,63). Indeed, research on stereotype threat shows that in performance contexts that do not support diversity, women and racial minority individuals show performance deficits (64). Furthermore, biomedical doctoral trainees from underrepresented groups, for example, often report being mentored unsatisfactorily, as their mentors may not care to understand their specific needs and challenges (1). The impact of unequal privilege and systemic barriers in organizational networks is pervasive as is evident in the historically excluded groups in biomedical and STEM arenas (14).

Generally, perceptions about CSE are formed throughout one’s academic career and can vary by gender and race (65,66); in fact, most research on the effect of gender and race CSE comes from K-12 schooling and undergraduate settings (e.g., 65,67). Thus, the CSE of trainees are shaped throughout their K-12 experiences, as well as undergraduate and graduate training experiences. With this knowledge, it follows that at the outset of career interventions, such as NIH BEST, personal trainee characteristics, such as gender and race, can also be expected to impact their CSE. There is a lack of clarity whether gender- and race-based differences in CSE in undergraduate training are extended into the biomedical graduate and postdoctoral training contexts. This knowledge gap might be rooted in the fact that, despite the challenges, some biomedical graduate students have indeed felt self-efficacious enough to choose STEM majors, despite the reputation of these fields are unwelcoming to underrepresented minorities (3, 23).

Individuals from underrepresented groups are also most at risk for attrition, and principal investigators (PIs) who serve as mentors can play an important role in nurturing graduate trainees’ CSE (37). Thus, understanding if trainees’ gender and racial identities differentially impact their CSE could help mentors and policy makers to address proactively systemic barriers to career persistence.

### Articulating a Blind Spot: The role of the intersectional perspective

As early as 1987, Lent and Hackett recommended focusing on the interrelated effects between race and gender when outlining the SCCT. Despite calls over the years (e.g., (68) and theorizing to date, research that takes an intersectional perspective remains sparse. Indeed, this blind spot has recently been highlighted again in Liu, Brown, and Sabat’s (21) call for scholars to focus their efforts on understanding the impact of intersectionality on core career constructs (21).

As described earlier in this section, research shows that individuals with multiple minority identities may be at a greater risk of facing emotional distress and poorer career outcomes than those who possess only one minority identity (63). This dilemma is captured in the double jeopardy hypothesis, also known as the additive model (69). This hypothesis suggests that patterns of discriminatory behavior are experienced as even worse if an individual inhabits two or more minority social identities. For example, in a study on women of color, Liu, Brown, and Sabat (21) found that compared to men and White women, women of color were more likely to be employed in two-year or four-year degree-granting institutions than in doctoral degree-granting universities that usually carry a higher status (21).

However, scholars suggest that there may also be a buffering effect of belonging to multiple minority identities, and that the negative impacts can neutralize each other. In a qualitative study of Black male professionals (doctors, engineers, among others), respondents noted how their intersectional identity also had positive effects (e.g., heightened social support from others with similar identity in the organization, and being held in high esteem for their competence and economic mobility (58)). In a similar vein, more experience with being a token representative can be less detrimental to minorities as they are more prepared to manage such devaluing interactions (70).

### Career Interest and Seniority in Training

Only more recently has scholarly work on graduate and postdoctoral education and training research shed light on factors influencing trainee career choice (e.g., (71,72); including intersectional identities, (2, 4, 23, 27, 73)). Furthermore, whereas self-efficacy is examined at various stages of education, there is a paucity of scholarly work that addresses self-efficacy change as a function of time spent within any particular training stage (e.g., seniority of trainee), particularly at the graduate or postdoctoral levels. While different stages of education such as undergraduate, post baccalaureate, graduate, postdoctoral levels are the focus in a few cases (74), such examinations generally concentrate on the related but non-synonymous construct of research self-efficacy (e.g., undergraduate, (75,76; graduate, (77); postdoctoral, (23), rather than CSE, as in the current work. Although some previous studies address career choice changes within the graduate level (2, 72, 78), they are often not combined with intersectional social identities, which are more frequently examined at the undergraduate level (67,68,79).

Furthermore, the authors are not aware of any studies that examine these variables (gender, race/ethnicity, career choice, and seniority) combined across graduate and postdoctoral training. The authors hypothesized that seniority and experience during training might impact CSE. Therefore, investigating an exploratory effect that may be moderated by intersectional social identities was proposed, such that these components may interact with gender and race/ethnicity.

### Research Aims

Taken together, current research suggests that one’s status as a member of an underrepresented minority group—specifically one’s gender and race—can impact one’s CSE. These findings also prompt the need to study these multiplicative effects in a biomedical context (including trainees), where diversification of the workforce is critically needed to maintain internal competitiveness. Therefore, the authors of the current study aimed to answer five primary questions using the data collected from the NIH BEST survey of trainees: (RQ1) What are the effects of *gender* on CSE? (RQ2) What are the effects of *race* on CSE? (RQ3) What are the effects of *interest in a PI career path* on CSE? (RQ4) What are the effects of *seniority within training stage* on CSE? and (RQ5) What are the multiplicative effects of *gender and race* combined with *career interest and seniority* on CSE?

## Materials and Methods

### Participants

All trainees eligible to participate in NIH BEST programs were invited to fill out extensive standardized surveys, irrespective of participation in career and professional development programming at their institutions (see Methods (25); see Supplemental Information S2 for sample questions). The sample consisted trainees predominantly in the biomedical sciences from 17 institutional sites in the United States participating in the NIH BEST awards (see (80) for program highlights).

Each institution either attained approval of its own institutional review board (IRB; e.g., Rutgers IRB#: E15-050; UNC IRB# 14–0544; Vanderbilt IRB# 190288) or relied on the NIH-approved data-sharing agreement (e.g., IRB Exemption Protocol ID#: 1412005184, OMB# 0925-0718) (25). Studies at each institution were generally considered either non-human subjects research or exempt, and as such approved in varying formats per institutional IRB requirements (e.g., an introductory survey statement/consent statement, information via email invitation, or a formal information sheet and/or consent form) that typically included information regarding the voluntary nature of participation, length of time for each component (e.g., survey estimated time), and/or intended purposes and limitations of data use.

As only US and naturalized citizens were included for analyses in this manuscript, all percentages hereafter are reported as the percentage of valid response totals within this defined sample. Temporary visa holders and green card holders were excluded from this analysis due to potential differences in cultures and residency status experiences. This international population will be analyzed separately in subsequent projects.

The current study analyzed common responses from standardized surveys across institutions, including demographic data. Trainee categories included graduate and postdoctoral trainees. Respondents self-identified their gender, race, and ethnicity. Options for gender included: male or female. No gender data were omitted in our sample. Female trainees are considered underrepresented by gender. Options for racial groups included: White, Asian, Black/African American, Native Hawaiian or Pacific Islander, and Native American.

Respondents also self-reported having Hispanic ethnicity (or not). Race or ethnicity was not disclosed by a subset of respondents (n = 270). Underrepresented trainee status by race or ethnicity (referred to as UR) was defined in accordance with NIH guidelines current at the time of survey design (25). Hence to protect participant identities and provide a robust sample size, a composite bivariate variable of race/ethnicity was created for the analysis, comprised of participants from well-represented (WR) or underrepresented (UR), with UR defined as identifying with one or more of the underrepresented racial or ethnic categories (Black/African American, Native Hawaiian or Pacific Islander, Native American, and/or Hispanic). The remaining responses that did not include one or more underrepresented identities were coded as WR (White, Asian, non-Hispanic).

Overall, 10803 responses were collected (of those, 6077 consisted of usable data for the proposed analyses). The current study sample included 6077 trainees who identified as US-born (n = 5541, 91%) and naturalized citizens (n = 536, 9%), including 71% graduate students (n=4283) and 30% postdoctoral trainees (n=1794). The sample consisted of White (n=4837, 82%), Asian (n=793, 13%), African American (n=369, 6%), Native American (n=92, <2%), and Hawaiian/Pacific Islander (n=36, <1%); with 91% identifying as non-Hispanic (n=5415) and 9% Hispanic (n=582). Respondents included 42% men, 59% women (see **Table 1**). Overall, 85% of respondents (see **Table 1**) were categorized as well-represented (White or Asian, non-Hispanic), whereas 15% were categorized as underrepresented in science (African American, Native American, and Hawaiian/Pacific Islander, or Hispanic).

**Table 1.** Number of respondents by race/ethnicity and gender.

| Intersectional Crosstabs |  |  |  |  |
| --- | --- | --- | --- | --- |
|  |  | Gender |  | Total |
|  |  | Men | Women |  |
| Race/Ethnicity | Well-Represented | 2119 | 2832 | 4951 |
|  | Underrepresented | 331 | 525 | 856 |
|  | Not reported | (122) | (148) | (270) |
|  | Total | 2450 | 3357 | 5807 |

### Data Collection

The current dataset included entrance surveys (i.e., surveys administered to trainees at the beginning of institutional participation in the NIH BEST-funded program) completed during the first and second years of the institutions’ NIH BEST funding (e.g., (25, 81,82)). Entrance surveys were chosen to ensure that the study population most closely mimicked that of the national population of biomedical doctoral and postdoctoral trainees. Entrance surveys also provided the most power and largest sample of trainees from UR populations. The questionnaire content and survey wording were developed by an external contractor (25) in response to requested topics identified by the NIH BEST Consortium.

### Measures

The entrance survey included five items that assessed CSE. Sample items included: Assess your abilities “to pursue your desired career path(s),” “to determine the steps needed to pursue your desired career path(s),” and “to seek advice from professionals in your desired career path(s)” (see Supplemental Information S2 for full list; and (82) for more information).

Respondents answered on a Likert scale from 1 = “Not at all confident” to 5 = “Completely confident.” Based on respondents’ answers, a composite score was created by summing total items, which were then averaged (mean of the 5 items) to create an Average CSE Score. Those participants who scored higher are assumed to have displayed a higher degree of self-efficacy than those who scored lower. Analyses were conducted using *mean* score. However, we found this difference noteworthy and is discussed further.

An expanded version of the CSE Scale (4-item version previously validated in (82) Cronbach’s alpha = .85; the current analysis retained all 5 items with similar reliability) met acceptable internal reliability criteria in this study sample (see Supplemental Information S3 for Chronbach’s Alpha analysis). Items of interest were assessed using Cronbach’s alpha to ensure they were related and provided reliable insight (Cronbach’s alpha = .86). Cronbach’s alpha is a measure of internal consistency or how closely items in a set are related to each other. Good internal consistency is indicated by a Cronbach’s alpha score greater than 0.70 to show high internal consistency, and that was exceeded in this study sample.

Year-of-training was also collected for both graduate and postdoctoral trainees, which was coded into a “junior vs. senior trainee” variable. In their fourth year and above, graduate students were considered senior (versus junior, third year and below); postdoctoral trainees were considered senior starting from the second year (versus in their first year of training). Graduate student seniority was based on typical milestones that are required by the third year of training, such as qualifying or comprehensive exams, advancement to candidacy, and completion of required coursework. Because postdoctoral training times vary substantially, postdoctoral seniority was defined simply as completing their first year. The expectations for a postdoctoral position may vary from lab to lab, but for this study, it was defined as such in consideration of the importance of the initial year to get established and set up a research plan, an experience qualitatively different from subsequent years. Therefore, the total number of years in a postdoctoral position was indicated as one total number regardless of whether more than one postdoctoral position was held.

Trainee career interests were recorded for 20 common biomedical career pathways, with response choices ranging from 0 (Not familiar enough to decide), 1 (Not at all considering), 2 (Slightly considering), 3 (Moderately considering), 4 (Strongly considering), to 5 (Will definitely pursue) for each career pathway. For the current study, the career pathway item “Principal investigator in a research-intensive institution” was used to determine each respondent’s interest in becoming a principal investigator. Responses of 4 or 5 were coded as 1, and responses of 0, 1, 2, 3 or non-response were coded as 0. For simplicity, hereafter the abbreviation for this variable will be referred to as “career interest” and response options referred to as “PI” and “non-PI”, respectively.

### Planned Analyses

The data were analyzed using between-subjects analyses, including a one-way ANOVA (trainee status) and a four-way factorial ANOVA (race x gender x career interest x seniority) using the NIH BEST entrance survey response data submitted by all participating universities. The current work proposed that intersectional effects of belonging to underrepresented groups and genders in the biomedical sciences may lead to disparities in CSE, and that career interests and seniority level may modify these disparities. All analyses were produced using SAS 7.15 HF9. Figures were completed in Graphpad Prism 9.1 and Illustrator 24.0.2. An Alpha of .05 was used to identify significant effects, with post-hoc Tukey corrections for multiple comparisons when relevant (two-tailed, between subjects t-tests). No outliers were removed. In cases in which some response data were missing for a composite variable but any responses were recorded, an average of the composite variable was still included in the analysis, adjusted for the total number of items recorded; in cases for which values for an entire variable were missing, missing data were automatically excluded from the analysis by the software. Note that the term effect is intended to refer to the empirical statistical relationship between variables throughout, and does not imply a causal relationship, but rather an observed association.

## Results

### Trainee Type

To control for any possible differences in experiences between different training populations, a preliminary analysis was conducted to assess whether there was a difference in reported CSE based on the training stage (predoctoral vs. postdoctoral trainee). Because the literature on CSE of biomedical graduate and postdoctoral trainees as a function of tenure in graduate school and beyond is sparse, the present study aimed to include an examination of these effects. Testing graduate versus postdoctoral training stage as a possible control variable revealed no statistically significant effects or interactions on CSE with the intersectional identity variables of interest (gender, race/ethnicity). Because the data analysis did not indicate any impact of training status on CSE, the remainder of the analyses were collapsed across the trainee populations. Hence, the central analysis consisted of a four-way factorial ANOVA examining the effects of race/ethnicity, gender, career interest, and seniority on the average self-efficacy beliefs across a combined sample of graduate and postdoctoral trainees.

### Gender, Race/Ethnicity, Career Interest, and Seniority

The four-way factorial analysis of variance (ANOVA) was conducted between gender (male and female), race/ethnicity (UR and WR), career interest (PI and non-PI), and seniority (junior and senior) on the level of CSE. The impacts of both social identity categories— gender (RQ1, p = 0.001) and race/ethnicity (RQ2, p = 0.001) —were each statistically significant with respect to mean CSE. As expected, based on previous research, displayed in **Figure 1a**, women were less self-efficacious (*M* _Women_ = 3.45) than men (*M* _Men_ = 3.54). As shown in **Figure 1b**, underrepresented groups were more self-efficacious (*M* _UR_ = 3.55) than well-represented groups (*M* _WR_ = 3.48). In addition, a trending interaction between Gender and Race/Ethnicity on CSE might suggest further investigation, though this conclusion should be interpreted with caution. This observation was identified as potentially important, but did not exceed statistical significance (RQ5, p=.05, see **Table 2**), and is further discussed as relevant in subsequent analyses and as related to significant two-way and three-way interactions. Yet the stratified effects of race/ethnicity may be more fully understood when examined across intersectional identities, career interest, and seniority (see interactions).

**Table 2.** 4-Way ANOVA: Race/ethnicity, gender, career interest, seniority. A four-way factorial analysis of variance of race/ethnicity x gender x career interest x seniority confirmed the two main effects from the original interaction, identified an additional main effect of career interest, and two significant interactions (career interest x seniority and race/ethnicity x gender x seniority). *P* values indicate level of significance, ****p<0.0001, ***p<0.001, **p<0.01, and *p<0.05. Find <u>interactive data visualizations</u> beyond Figured 1 & 2 provided via Tableau (see Supplemental Information S4 for more information).

| ANALYSIS OF VARIANCE |  |  |
| --- | --- | --- |
| Variable (s) | *F-Test* | *P-Value* |
| Race/Ethnicity* | 12.27 | <.001 |
| Gender** | 10.90 | <.001 |
| Career Interest*** | 80.16 | <.0001 |
| Seniority | 3.02 | 0.08 |
| **Race/Ethnicity x Gender** | **3.83** | **0.05** |
| Gender x Career Interest | 0.21 | 0.65 |
| Race/Ethnicity x Career Interest | 0.85 | 0.36 |
| **Race/Ethnicity x Seniority\*** | **5.04** | **0.02** |
| Gender x Seniority | 0.84 | 0.36 |
| **Career Interest x Seniority\*** | **6.14** | **0.01** |
| Race/Ethnicity x Gender x Career Interest | 0.06 | 0.80 |
| Race/Ethnicity x Gender x Seniority | 0.01 | 0.93 |
| **Race/Ethnicity x Career Interest x Seniority\*** | **5.00** | **0.03** |
| Gender x Career Interest x Seniority | 0.63 | 0.43 |
| Race/Ethnicity x Gender x Career Interest x Seniority | 1.27 | 0.26 |

**Figure 1.**
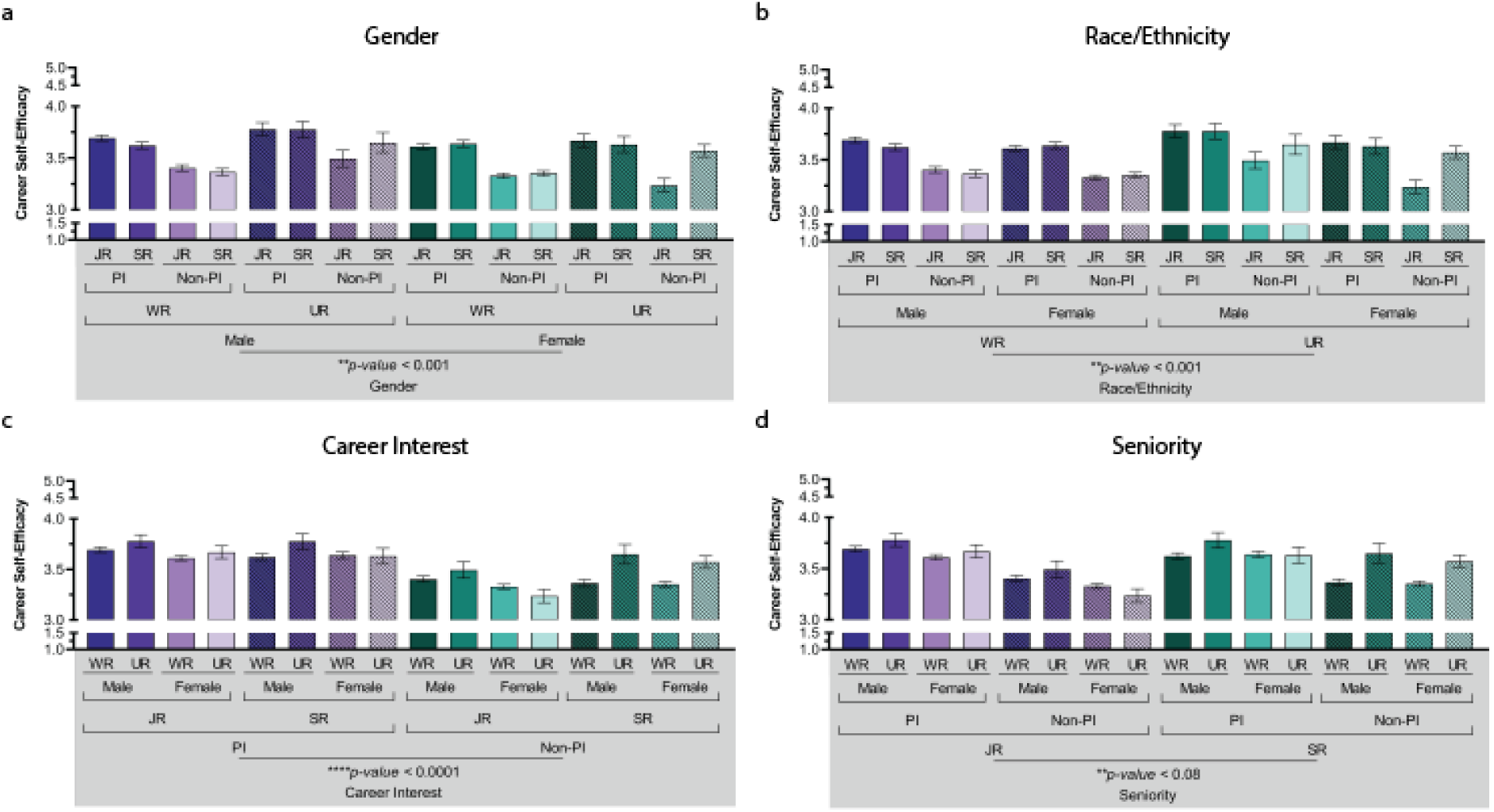
Main effects of gender, race/ethnicity, career interest, and seniority Main effects of gender, race/ethnicity, career interest, and seniority. All main variables showed a significant effect on self-career efficacy, with the exception of seniority, which was trending but not significant. A four-way ANOVA of gender, race/ethnicity, career interest, and seniority on the level of CSE showed significant main effects for each variable. P-values indicate the main effects for each variable of interest, ****p<0.0001, **p<0.01, and *p<0.05. (WR=Well-Represented, UR=Underrepresented, JR=Junior, SR=Senior). Color differences (green and purple) indicate main effect for each variable. For 1a gender (male vs. female), 1b race/ethnicity (WR vs. UR), 1c career interest (PI vs non-PI), and 1d seniority (JR vs. SR).

Main effects were detected between PI versus non-PI for career interests among trainees (RQ3, p < 0.0001), such that trainees with interest in working as a PI at a research-intensive institution were more self-efficacious (*M* _PI_ = 3.65, *M* _Non-PI_ = 3.37) displayed **in Figure 1c**. Only marginal main effects were identified between senior compared with junior trainees, but did not reach significance (RQ4, p < 0.08); nonetheless, a trend of senior trainees expressing more CSE compared to junior trainees may still be of interest (*M* _Junior_ = 3.49, *M* _Senior_ = 3.49). Significant interactions (RQ5) included 2-way interactions between seniority x race/ethnicity (*p* = .02) and seniority x career interest (*p* = .01), along with a 3-way interaction between race/ethnicity x career interest x seniority (*p* = .03; see **Table 2** and **Figure 2**).

**Figure 2.**
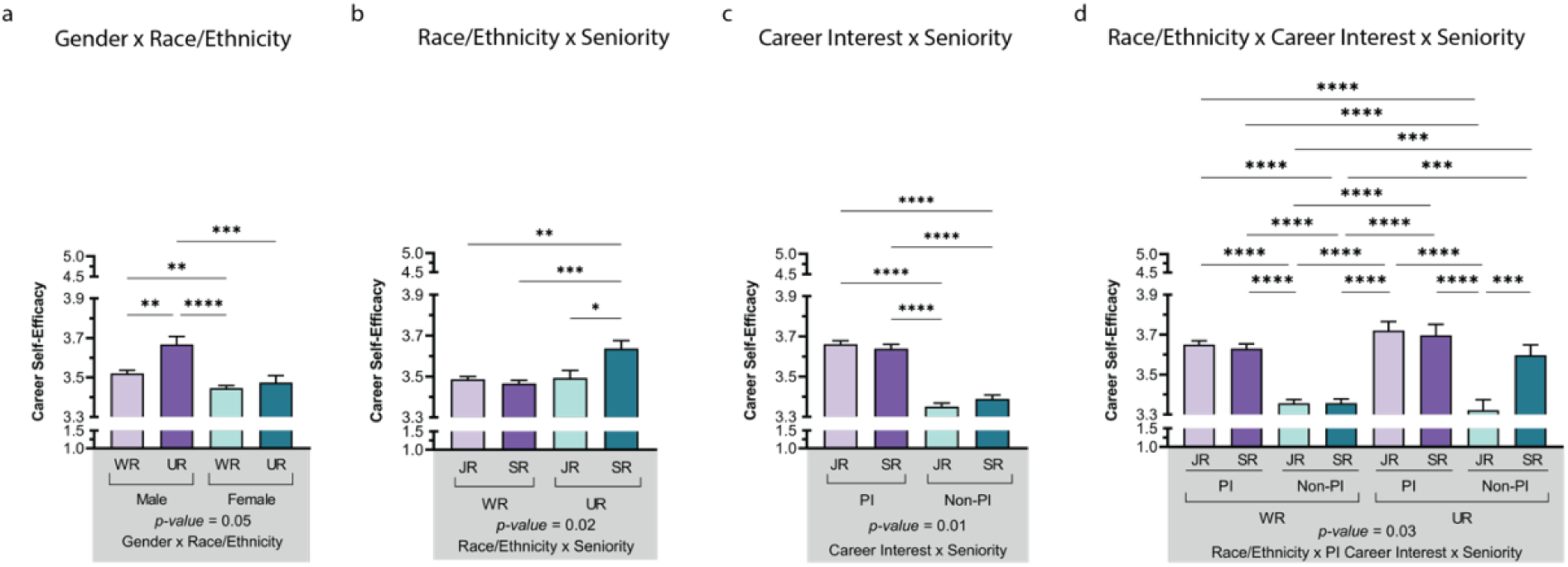
Interactions of gender, race/ethnicity, career interest, and seniority. Patterns for gender and race/ethnicity were generally similar to the primary analysis; however, they also showed interactions with career interest and seniority. Career interest showed a significant main effect, with higher CSE for those strongly interested in becoming a PI at a research-intensive institution compared to those with moderate -to-no interest in this career path. The effects of seniority trended slightly higher for junior trainees than senior trainees, but differences were more pronounced in the career interest x seniority interaction, which attained significance. Post-hoc t-tests were conducted between all possible pairings as illustrated by each end of the respective bracket. P-values indicate significance of Tukey’s multiple comparison tests (see Supplemental Information S5 for full table of values), ****p<0.0001, ***p<0.001, **p<0.01, and *p<0.05. Significance values at the bottom of each panel (a-d) indicate interactions. Panel 2d displays the 3-way ANOVA with only multiple comparisons not represented in other panels included (e.g., not displayed in panels 2a-c). *Note:* WR=Well-Represented, UR=Underrepresented, JR=Junior, SR=Senior. Color differences (green and purple) indicate the primary variables. For 2a gender (male vs. female), 2b race/ethnicity (WR vs. UR), 2c career interest (PI vs. non-PI), and 1d race/ethnicity (WR vs. UR) and career interest.

Although gender (**Figure 1a**) and race/ethnicity (**Figure 1b**) showed significant main effects, and seniority showed a trending main effect (**Figure 1d**), the most robust effect was seen for career interest (see **Figure 1c**). Interactions between race/ethnicity, seniority, and career interest also showed some interesting combinations. Of note, men belonging to UR groups had higher CSE than women belonging to either WR or UR groups (no different from WR men; see **Figure 2a**). Senior trainees belonging to UR groups had the highest CSE compared with all other groups (**Figure 2b**). While career interest (PI) had a positive relationship with CSE, senior trainees interested in PI careers were rated highest in CSE compared with other combinations (**Figure 2c**). Finally, senior WR and UR trainees who were interested in a PI career were statistically more career self-efficacious than most other combinations of junior/non-PI–with the notable exception of trainees from UR groups who were not strongly considering becoming a PI (senior/UR/non-PI; see **Figure 2d)**. Of note, males from UR groups interested in a PI career had noticeably higher CSE than other groups (Male/UR/PI *M* = 3.78, both junior and senior; see **Table 3**). Hence, those with the highest group means included UR/male/PI, which was higher than other intersectional combinations, though other senior/PI groups were next highest (ranging from *M* = 3.62-3.78; see **Table 3**).

**Table 3.**
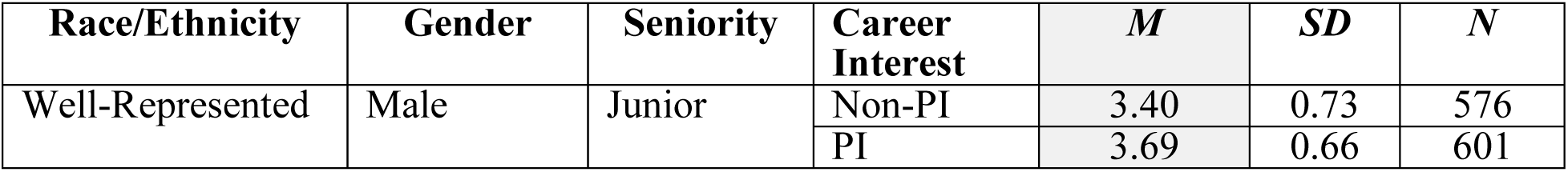
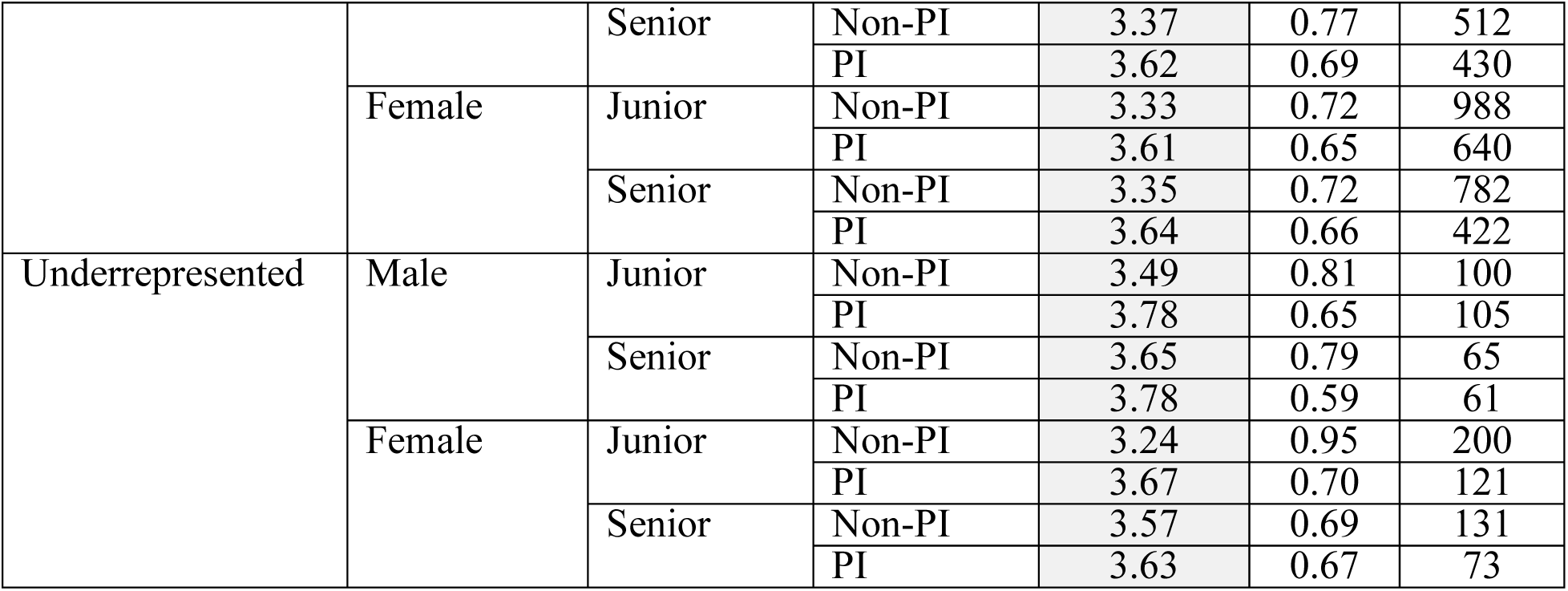
Patterns of Race/Ethnicity, Gender, Career Interest, and Seniority. Patterns of race/ethnicity and gender were generally maintained, with newly identified interactions with seniority and career interest. Of note, UR junior trainees strongly considering a PI career were particularly high in CSE (male and female), as were UR senior male trainees interested in becoming a PI (see bold). *Note:* Due to some respondents skipping questions, the N above represents the population that were included in the analysis.

## Discussion

CSE, defined as one’s beliefs in the ability to be successful in a specific career, is a key predictor of career intentions (83,84). One of the main goals of the research study presented here was to assess how gender and race/ethnicity relate to CSE of graduate and postdoctoral trainees. Our finding that women reported lower CSE than men is in line with previous research (3, 23, 85). However, in addition to studying the effects of gender and race/ethnicity separately, the current study posits that it was vital to take an intersectional identity perspective to understand the multiplicative effect - of gender and race/ethnicity combined - on trainees’ CSE. Building upon prior work that had validated the CSE in the NIH BEST context (82), the current work extends findings for CSE by including intersectional identities. In the NIH BEST national survey sample, men and women from underrepresented racial/ethnic groups reported higher CSE than men and women from well-represented groups. Furthermore, these findings show that belonging to two intersecting identities (i.e., being female *and* being a racial/ethnic minority) may offer a multiple identity advantage (as defined by (10)) to CSE in the biomedical sciences – but with some nuance. Senior female trainees from UR groups, interested in both PI-and Non-PI careers showed higher CSE (as well as those more junior who were interested in PI careers). However, junior female trainees from UR groups who were not interested in PI careers showed the lowest intersectional identities across all groups. It is important to investigate the finding that there is some evidence of a multiple identity advantage, especially given that existing literature shows that individuals from underrepresented groups tend to report lower CSE than well-represented individuals (86). In another study, postdoctoral trainees from underrepresented groups versus postdoctoral trainees from well-represented groups showed no significant differences in outcome expectations related to CSE but did show higher research self-efficacy (23). Perhaps the large national sample size of the current study (n=6077, power for detecting omnibus fixed effects, specials, main effects, interactions = 95%; calculated by G*Power) provided a greater power to detect effects that may have been obscured otherwise. Power to detect main effects was generally quite good (Gender, PI Career, and Seniority = 99%; calculated by NQuery), but was insufficient for finding effects for race/ethnicity due to the still limited sample sizes (power = 28.5%; calculated by NQuery). Accordingly, even with this comparatively large sample, the power to detect some UR and WR differences could still have been obscured, hence the recommendation to revisit the trending interaction with race/ethnicity and gender in future studies. The results may help clarify some of the mixed outcomes of prior studies related to effects of gender and race/ethnicity that are likely to have been hidden due to the underpowered nature, and the lack of using an intersectional approach (2,4). Additionally, such a disciplinary focus on biological and biomedical trainees may show distinct patterns from other fields of study.

Interest in pursuing an academic PI career showed complex relationships with key variables of seniority and race/ethnicity. Generally, CSE for those strongly considering a career as principal investigator at a research-intensive university (PI) was higher compared with those not strongly considering that career path (Non-PI); yet these relationships were also dependent upon seniority (all three interactions), as well as race/ethnicity (two interactions).

Historically underrepresented gender and race/ethnicity were associated with lower CSE (main effects), yet the interaction was not significant (two-way interaction); since it was trending, however, this observation may be worth future investigation due to the marginal significance of the interaction. The relationship between career interest (considering a career as a PI at a research-intensive university versus those not considering this career pathway) and CSE was a strong predictor. Those respondents who actively expressed interest in becoming a PI at a research-intensive institution (PI) showed greater CSE compared with those not (non-PI). In contrast, Lambert and colleagues (24), found that the most productive postdoctoral trainees (e.g., more publications) with a higher research self-efficacy, a related construct, were choosing to leave academia (23).

Although trainee-type (i.e., graduate versus postdoctoral trainees) did not show significant differences, seniority (within each training stage) did show significant effects. Moreover, seniority and career interest combined to predict unique patterns (two-way interaction), as did race/ethnicity, career interest, and seniority (three-way interaction). Career theories (87) suggest that accruing experience in one’s domains of learning builds higher CSE as individuals become adept at responding to the demands of their career (88,89). Based on this rationale, the current study anticipated that advanced doctoral students would show higher CSE than entry-level graduate students. The data show that in the biomedical context, spending additional years in training positions was indeed related to higher CSE. However, there were some interactions with seniority, such that junior trainees interested in PI Careers showed especially high CSE. It is likely that trainees, as they gain experience in their field, learn that the path to finding a job that fits their own values and needs is not as straightforward as they may have anticipated early in their careers, especially in careers to which they had limited exposure. This outcome may suggest that less confidence could be addressed by early career exploration and training activities, particularly for those interested in non-academic career pathways; however, with lower than anticipated CSE across the board in this sample, building self-efficacy at any stage would provide a valuable training addition. Future research should continue to examine the impact that career progression and experience have on CSE.

To complement the extensive work addressing such questions at the undergraduate research experience, new investigations could include research on the process of gaining CSE during training and who might be impacted differentially during the training years, along with how and why CSE is acquired. Such results would be particularly relevant to better inform the development of program support strategies for junior doctoral trainees, who appear to be most in need of such interventions in comparison to more senior trainees. One possible explanation is that each time trainees transition to different stages, many of those who choose pathways other than the academic route may matriculate out into the workforce, leaving behind those who are still considering academic career pathways, such as becoming a PI, to continue in the postdoctoral training track (e.g., self-selecting those who remain into a more confident PI-career-focused cohort). Of course, lab environment, institutional environment, and personal interactions and especially mentoring experiences could impact the development of CSE and/or the exposure to and selection of career pathways.

Junior trainees displayed lower CSE than senior trainees suggesting that it may be built over time, which is encouraging. However, worthy of note, group means for CSE ranged only from 3.11 to 3.80, which is much lower than might be expected at these advanced stages of training where one might expect to see ranges more in the 4-5 ranges based on comparable studies in related fields). The overall low CSE of advanced trainees may be indicative of a larger issue around the scientific training process, exposing trainees to repeated failures, hypotheses, which could not be verified, experiments that need to be redesigned or repeated, and minimization of success. It may be that faculty mentors are so focused on robust analyses and critiques that they may fail to routinely recognize early-career scientists’ accomplishments with positive reinforcement. The process of positive enforcement to reward errors in training, also known as Error Management Training, has been shown to lead to better training outcomes, across a meta-analysis of a broad range of studies (90). To date, faculty often have limited or no formal training in good mentoring practices or knowledge of the myriad of careers for which their protégé’s’ skills are useful; increasing faculty knowledge of practices and resources related to productive mentoring might help to enhance their graduate and postdoctoral trainees’ CSE as they transition to graduate and postdoctoral training.

### Implications

The low CSE of advanced degree scientists is somewhat concerning. Finding systemic ways of increasing opportunities to build trainee CSE, through mentor-training and direct training and reinforcement for graduate and postdoctoral trainees, is important for future considerations. Evidence-based mentoring principles (developed through research at the Center for the Improvement of Mentored Experiences in Research, CIMER) point to important factors that can especially influence self-efficacy (the four primary SCCT factors, with intersectional differences by population; e.g., (91)). The role of mentoring may be vitally important for trainee persistence towards a specific goal. Intervention programming, for mentors (e.g., CIMER) as well as direct delivery particularly to UR trainees (for undergraduates: Maximizing Access to Research Careers, MARC; Research Training Initiative for Student Enhancement, RISE; Stipend for Training Aspiring Researchers, STAR; McNair Scholars Program; and inclusive of higher degree programs such as the Initiative for Maximizing Student Development), could truly make a difference in confidence-building and the way exposure to other fields is handled that will protect confidence (including potential exposure to other academic and non-academic careers).

Mentoring conversations and role models to build self-efficacy may be particularly crucial at early stages of training (i.e., for junior trainees). Other more longitudinal studies have examined the declining interest in an academic career over the course of PhD training (78). Exposure to the reality and attainability of their academic career goals influences students’ perceptions of their own research abilities. Roach and Sauermann (78) find that early in their graduate training students are not familiar with diverse non-academic career options that are available to them and gaining this information as they progress with their graduate training does play a role in their decline in interest in an academic career (78). Considering different events and realizations that may occur during transition periods from junior to senior status may include the research environment and even specific events that may affect self-efficacy. For example, passing the PhD candidacy exam (typically second or third year) could be a turning point where students start to look beyond their original goals. Similarly, for postdoctoral trainees, an influential factor could be getting established in one’s lab, gaining familiarity with institutional and laboratory support levels and resources, building a relationship with the research advisor/PI, and evaluating the potential for high impact publication(s) for postdoctoral research project(s). Other possible factors may include realizing the full spectrum of one’s research expertise, recent history of independent project/funding troubles or successes (8), or exposure to a potential dream job or career. In other words, there may be specific events in graduate and postdoctoral training that could contribute to differing junior and senior trainee experiences. Hence the central question may be, *how can mentors, programs, and institutions better support trainee CSE during these crucial stages of development and across the training period?*

Furthermore, previous studies have identified differences in values impacting career choice amongst women and trainees from UR groups who might otherwise choose to stay in academia (4, 23, 73). These differences are reflected in the current study results by the comparatively high CSE evidenced by senior trainees from UR groups who are interested in non-PI careers, presumably choosing to exit the academic faculty career pathway. UR and women trainees value giving back to the community and university through mentoring, teaching, and community outreach – all of which are undervalued by the current research enterprise and are at best uncompensated and unrecognized work, which is actively discouraged for faculty promotion purposes. This indicates a need for a fundamental transformation of the academic culture and workforce roles, if it indeed purports to recruit and retain diverse faculty participation. In addition, other factors may play a role in career choice such as variety, prestige, and salary, as well as family influence, PI mentoring or encouragement (4). These factors are further evidence that transformational change in academic culture and compensation is much needed, including addressing academic pay inequities and incentives to attract and retain more women and candidates from UR groups; valuing contributions of faculty through financial compensation, promotion, and tenure across a wider variety of activities, including mentorship and community outreach, and improving outcome expectations for careers. Fundamentally changing academia to be more inclusive, expanding value and recognition of contributions, and creating funding equity, all rely on the role of mentors and PIs in taking sustainable, collective action (92).

### Limitations & Future Directions

The current study focused on a domestic sample, and the findings cannot be generalized for international trainee populations who occupy intersectional identities by virtue of their gender, race/ethnicity, culture plus their status as immigrants in the US, thus facing a host of additional barriers and difficulties, However, because biomedical fields attract a sizeable population of international trainees (over the preceding decade, nearly 40% of PhD earners in science and engineering were awarded to temporary visa holders; (93), they are clearly an important subgroup to study.

The analysis in our study was limited by gender options (male, female) offered in our survey, whereas a non-binary gender selection was not available. This restriction limits our ability to account for gender non-binary persons in the current sample, and serves as a potential area that could be expanded to include other groups in the definition of being underrepresented in the sciences. This population of trainees is highly understudied, although non-binary trainees represent this group contributes significant and valuable work to scientific fields (95). Especially given association between gender and CSE, it will be important for future studies to examine patterns among gender non-binary trainees. Furthermore, as with other historically underrepresented groups in science, the inability to achieve large sample of respondents in research studies has limited their inclusion in research, making it important for future studies to recruit robust and representative samples of gender diverse individuals. Findings reported here may not be generalizable to a larger, more inclusive group of trainees.

The present study also provides analyses on a limited number of UR individuals, categorized on the basis of race/ethnicity and gender, compared to WR individuals. Nonetheless, a potential reason the number of UR persons in the sample was lower than our WR group, may be partially due to the number of UR that decide to leave or not pursue graduate programs in STEM altogether (94). The loss of such talent creates a significant gap in our knowledge of the intersectionality of race/ethnicity and gender on career efficacy.

Given the attrition issues seen in STEM contexts especially the higher rates for UR individuals (94), it is vital to note that our results demonstrating some benefits for UR individuals are viewed with caution. Specifically, higher CSE in UR (senior) versus WR individuals could simply stem from the fact that the sample in our study is comprised of UR individuals who are still in STEM doctoral and postdoctoral careers (i.e., survivorship bias). They may have had to survive far greater odds to be at this career stage than WR trainees, so they report higher CSE. However, they are not the full population of UR by any means and while this would be a reasonable alternative explanation for some of our findings, it is important to note that junior female trainees from UR groups who were not interested in PI careers showed the lowest CSE of all groups. This sheds some doubt on survivorship bias as an alternative explanation, as even within some people of the current sample of “survivors”, there is lower efficacy observed. This observation lends support to our idea that it is the combination of career interest, seniority, gender and race that are implicated in an intersectional manner in these findings. Overall, it will be important for replications and experimental studies to further assess the multiple identity advantage hypothesis.

Another limitation is potential bias because the study was observational in nature and not a randomized controlled experimental selection process for participation in career development activities. Furthermore, while the entrance data were collected near the beginning of the NIH BEST awards, it is possible that NIH BEST selected institutions somehow systematically differed from non-selected institutions, such as having institutional support to apply for the grant or some piloted or existing professional development programs. Yet this seems an unlikely explanation, as a broad array of schools with differing levels of prior experience, including programs with and without an existing professional development infrastructure, were selected by the NIH.

Future directions might include testing the effects of interventional programs to increase earlier building of CSE for the pursuit of a PI career, as well as earlier building of CSE for discovery and preparation toward other career pathways of best fit. In the current analysis, the authors investigated the impact of existing individual differences on CSE, but for programs whose goals include training future PIs (e.g., CIRTL network), a strong self-efficacy-building component may be an effective strategy to increase such a career selection – particularly for junior women trainees, and junior women trainees from UR groups. It is possible that trainees may change their career choices throughout their training independently of their CSE, in which case an interest in a PI career and associations with self-efficacy may be incidental. However, given that the career interest had by far the largest effect size, it seems reasonable to interpret this strong relationship between CSE and career interest as related constructs. Future work should continue to explore this relationship and possible directions of causality to better understand how these variables may impact each other over time. Furthermore, the importance of providing programming and opportunities for career exploration and development to all trainees cannot be overstated. Yet, specific barriers faced by individuals from UR groups and gender identities still need to be specifically acknowledged and addressed, to include the explicit development of self-efficacy.

For future survey studies, strategies should be employed in order to increase the recruitment and retention of UR persons in advanced training programs, in addition to seeking UR respondents to increase representation of UR respondents in research, to ensure that we are fully encompassing the impact of intersectionality on CSE. Evidence-based strategies that can be implemented in NIH BEST affiliated institutions to mitigate the significant differences in UR versus WR persons have begun to be explored in prior work (95), yet more is needed. Such initiatives include the overall cultural shifts in the academic workplace towards institutes that support women of color faculty members. Other strategies specifically involved increasing access for women of color, as well as dismantling unconscious gender and bias through activities and various programs. While these strategies will not immediately solve the impact of systemic racism in academia, it will begin to address these issues at institutes that may not already be working towards a more inclusive academic atmosphere. If we can provide increased cultural awareness and representation in academic institutions, this would be a step in the right direction for increasing the number of UR individuals in STEM, as well as to increase representation in survey results, to better understand the effect of intersectionality on CSE.

Transitional programs, from undergraduate to graduate education programming (early career interventions; e.g., Post baccalaureate Research Education Programs or the Initiative for Maximizing Student Development), later training stage interventions (postdoctoral mentored research and teaching experiences; e.g., Institutional Research and Academic Career Award Programs) could constitute some of these intervention programs, as well as those interventions that take a talent development, mentoring, or coaching approach (e.g., talent development approach; (74); mentorship training, (91,96); career coaching, (27) that could contribute to building self-efficacy within trainees. Program curricula that are centered on support for trainees, including exposure to careers outside of academia and multiple aspects of academia outside of research (e.g., NIH BEST programming), and reinforcing trainee development beyond being a researcher, but also mentor, mentee, and teacher in expanded roles will collectively enhance the experiences for all trainees. Prolonged mentoring/interventions (31, 76, 97-99) could promote self-efficacy in academia, especially for training and exposure into specific careers in and beyond academia. For example, for those interested in becoming a faculty member or PI at a research-intensive institution, provide opportunities to explore related faculty and administrative career pathways at other types of institutions, such as teaching-intensive institutions (including primarily undergraduate institutions, small liberal arts colleges, historically black colleges and universities, minority-serving institutions). In addition, trainee-centered programming designed to increase awareness of the many career paths outside of academia to which their skills could be applied effectively to improve society broadly could also promote CSE for these additional roles based on fit. Additionally, future studies investigating whether CSE predicts career outcomes would translate the importance of studies such as ours to truly determine the impact of developing CSE of trainees across various training stages.

While these recommendations are relevant to all trainees, they are particularly important for UR trainees because of previous findings on the importance of different career values for UR scientists.

## Conclusions

Social identity was associated with higher CSE for individuals from underrepresented versus well-represented groups. Women expressed lower CSE compared with men. A trending interaction between race/ethnicity and gender provided partial support for the hypothesis that intersectionality impacts CSE, as did additional two- and three-way interactions with race/ethnicity, career interest, and seniority. Of note, women trainees generally show lower CSE, especially junior women trainees. Most impacted in this regard were junior women trainees from UR groups (whereas senior women from UR groups, especially on the PI career track, showed particularly high CSE). Further analyses revealed a trend of increased CSE between senior versus junior trainees, though CSE was lower than expected across the board, suggesting the importance of reinforcing career development opportunities and experiences for trainees. By far the largest effect was demonstrated related to interest in a career as PI at a research-intensive institution, with those expressing that specific interest having higher CSE; however, pattern was not evident senior trainees from UR groups who were not interested in a PI faculty career path. Interestingly, this group did not differ significantly from the two highest groups, senior trainees interested in a PI career from WR or UR groups.

Together these findings underscore the need for the transformation of the biomedical training and research communities in academia in order to develop and retain a diverse talent pool. Further studies are needed to extend these findings and implications for actionable policy and suggest the need for more research in this area. Despite the comparatively large sample of this study, the smaller sample size for trainees from underrepresented racial/ethnic groups nonetheless limited findings and analyses. Additional systematic evaluation and recruitment of trainees from underrepresented racial/ethnic groups in graduate education research is crucial to advance the field.

## Supplemental Information Files

(Available via Open Science Framework DOI: 10.17605/OSF.IO/4F8V5)

S1. Abbreviations

S2. Questionnaire items

S3. Inter-reliability of items

S4. Interactive data set availability

S5. Tukey’s corrected multiple comparisons

